# Olfactory responses of *Drosophila suzukii* parasitoids to chemical cues from SWD-infested fruit

**DOI:** 10.1101/2022.09.08.507209

**Authors:** F. Triñanes, A. González, G. J. de la Vega

## Abstract

Since *Drosophila suzukii* (Diptera: Drosophilidae; SWD) became a worldwide pest of soft-skinned fruits, multiple mitigation strategies alternative to insecticides have been explored. Among these, the search for biological control agents has prompted the assessment of drosophilid parasitoids for SWD control. Olfactometer bioassays with drosophilid parasitoids have shown that host substrate-related complex olfactory cues are relevant during host search. No information is available, however, on which fruit volatiles may be used as host-related cues. Here, we used gas chromatography coupled to electroantennography (GC-EAD) to evaluate the antennal detection of ecologically relevant fruit odours by two drosophillid parasitoids, *Leptopilina boulardi* (Hymenoptera: Figitidae) and *Trichopria anastrephae* (Hymenoptera: Diapriidae). We found that females of both wasp species are capable of detecting the main volatile compounds emitted by SWD-infested strawberries, five and ten days after oviposition by SWD females. The EAD-active fruit compounds were identified by GC-MS analysis as the common fruit esters ethyl butanoate, methyl hexanoate and ethyl hexanoate. The relative proportions of these fruit esters vary over time, with potential ecological significant for larval and pupal parasitoids. Our study is the first to report GC-EAD responses of microhymenopteran wasps of drosophilid flies. Understanding the sensory ecology of host-related chemical cues may be useful to optimize the biological control of *D. suzukii* by parasitoid wasps.

## Introduction

Plants structure and support multitrophic interactions in nature. Among other aspects, plants are conspicuous in their contribution to the chemical landscape of ecosystems. Plant phenological changes under herbivory are often accompanied by changes in associated volatile chemistry, changes that can be exploited by both herbivorous insects and natural enemies. Changes in volatile chemistry may provide cues about the suitability of a given substrate, as well as spatial and temporal information of a potential feeding or oviposition resource (De Moraes et al., 1998). Parasitoids exploit volatile semiochemicals from plants to find their insect host (Lewis & Martin, 1990; Vet & Dicke, 1992). Indeed, the detection of specific compounds may be an essential prerequisite for modulating host-searching behavioural responses, within the complexity of environmental and internal inputs (Anton et al., 2016).

A frugivorous species that gained worldwide notoriety for its negative impact on soft-skinned fruits is *Drosophila suzukii* (Diptera: Drosophilidae), also known as the spotted wing drosophila (SWD). Females use ripening berries and cherries as oviposition substrate, damaging the fruits close to their harvest (Walsh et al., 2011). The short time window between damage and consumption limits the use of insecticides for SWD control, highlighting the need for alternative management strategies. In the context of biological control and integrated pest management, the search for effective parasitic wasps against SWD received significant attention (Wang et al., 2020). Drosophilid parasitoids evaluated as SWD controllers have included larval parasitoids from the genera *Leptopilina* or *Ganaspis* (Hymenoptera: Figitidae), as well as pupal parasitoids from the genera *Trichopria* (Hymenoptera: Diapriidae) and *Pachycrepoideus* (Hymenoptera: Pteromalidae) (Rossi-Stacconi et al., 2015, Daane et al., 2016, Ibouh et al., 2019, Lee et al., 2019). SWD parasitoids need to find SWD-infested fruit and presumably exploit fruit volatiles as cues, a tritrophic interaction that is not well understood and may be key for successful biological control. The study of these interactions have included the behavioural assessment of wasps in olfactometer tests in response to natural volatile blends from infested fruit (Biondi et al., 2021; de la Vega et al., 2021; Wolf et al., 2020). Because these are complex and dynamic volatile blends, separating its components and evaluating their ecological significance may be useful to further characterize these interactions.

Coupled gas chromatography/electroantennographic detection (GC-EAD) is a widely used technique for identifying specific insect olfactory stimulants in complex volatile organic compound (VOC) blends. Briefly, the insect antenna acts as a selective biological detector in parallel with the output obtained from the normal GC detector, usually a flame ionization detector (FID) (Sullivan & Slone, 2007). Multiple studies have evaluated GC-EAD responses of braconid parasitoids of true fruit flies (Tephritidae) to host-related cues (Benelli et al., 2013; Ngumbi et al., 2009). To our knowledge, however, no previous studies using GC-EAD have been conducted with drosophilid parasitoids such as *Trichopria, Leptopilina* or *Ganaspis* species. Uncoupled electroantennogram (EAG) studies were conducted in *Leptopilina heterotoma* (Hymenoptera: Figitidae) (Vet et al., 1990), mostly focusing on the effect of wasp pre-exposition to host food odours on the wasp’s EAG response.

Here we report the use of GC-EAD to investigate the antennal responses of *Leptopilina boulardi* (Hymenoptera: Figitidae) and *Trichopria anastrephae* (Hymenoptera: Diapriidae) to VOCs from SWD infested strawberries. These two drosophilid parasitoids attack larvae and pupae, respectively, so we performed our experiments at five and ten days after SWD oviposition to account for the presence of larvae or pupae inside the fruit (Tochen et al., 2014).

## Materials and Methods

### Parasitoids

Adult wasps of *T. anastrephae* and *L. boulardi* were obtained from *Drosophila melanogaster* Meigen (Diptera: Drosophilidae) breeding tubes maintained on artificial diet (500 mL distilled water, 50 g glucose, 20 g bread yeast, 4 g agar, 40 g corn-flour, 1.5 mL propionic acid, 3.5 mL nipagin) and kept under incubator conditions (21.5 ± 1°C, 65 ± 5% relative humidity, 12:12 h photoperiod). Adult wasps were kept with access to a diet based on distilled water and honey (50:50) for 5 to 10 days prior to their use in experiments.

### SWD oviposition and VOC collection

SWD oviposition and VOC collection were performed in incubators set at 60% HR, 14:10 h D:L photoperiod and temperatures of 15 °C in darkness and 23 °C during daylight. Single clone strawberry plants were grown in pots under greenhouse conditions and transported to the laboratory with ripening strawberries. Individual strawberries still attached to their plants were enclosed with three mated SWD females during 24 h for oviposition. Five and 10 days after SWD infestation strawberries were individually enclosed in polyester oven bags (20 x 15 cm) attached to the peduncle with a plastic seal. An activated carbon filter was attached to the oven bag for incoming air, and a folded acetate sheet was placed inside the bag and around the fruit to prevent the bag from collapsing due the pump suction. Air with the fruit VOCs passed through a glass Pasteur pipette with 60 mg of HayeSep Q as adsorbent material, then suctioned by a portable pump (Casella, Apex2) set at 0.3 L/min. Retained compounds were desorbed with 1 mL of hexane, then 100 μL of a solution of n-tridecane was added as internal standard and the mixture concentrated to 100 μL under N_2_ for GC-MS and GC-EAD analyses. After the second VOC collection, 10 days after oviposition, SWD infestation was confirmed and quantified by carefully immersing and mashing the fruit in a saturated sugar solution, according to Dreves et al. (2014).

### GC-EAD analysis

Wasp antennae were removed with dissecting scissors, and the apical flagellomere and the scape were severed. The electric circuit consisted of two silver (Ag/AgCl) electrodes immersed in Beadle-Ephrussi Ringer solution (NaCl 128 mM, KCl 4,7 mM and CaCl_2_.2H_2_0 1,9 mM) inside microcapillaries; the signal electrode was pre-amplified in a Syntech combi-probe (10x) and further amplified by a Syntech amplifier (IDAC-2). The GC system was a Hewlett-Packard gas chromatograph (5890 series II) equipped with a DB-5 column (30 m x 0.25 mm x 0.25 mm, Alltech, USA) and a flame ionization detector (FID). A Syntech Stimulus Controller (Model CS-55) delivered humidified air to durect the volatiles eluted by from the GC towards the antennal preparation, with a continuous flow (1.05 L/min). FID and EAD signals were integrated using Syntech’s GC-EAD software (v.2014).

At least 10 replicates of VOC collections were obtained and analyzed 5 and 10 days after SWD oviposition. Of these, five representative samples of each VOC collection time were mixed in order to use an homogenous stimulus blend for GC-EAD replicates. For each GC-EAD run, one microliter of the VOC blend solution was injected in splitless mode with H_2_ as gas carrier (2 mL/min). The oven temperature started at 40 °C for 1 min, increased to 150 °C at a rate of 5 °C/min and to 250 °C at 10 °C/min (held for 1 min). Injector and detector temperature were kept at 250 °C, and the EAD interface temperature at 220 °C (Syntech TC-02). The column effluent was split with a 1:1 ratio inside the GC oven, using nitrogen (20 mL/min) as additional make-up gas before the column spliter. VOC extracts from each post-infestation time (5 and 10 days) were analyzed with 20 independent antennae from each parasitoid species, using left and right antennae equally.

### Chemical identification

Compounds that elicited an antennal response were identified by gas chromatography-mass spectrometry (GC-MS) using the same chromatographic conditions as detailed above on a QP5050 Shimadzu GC-MS equipment. Identification of the compounds was based on EI-MS fragmentation patterns using the NIST 17 database run on a GC-MS solution software (Version 4.45 SP1).

## Results and Discussion

The antennae of female *L. boulardi* and *T. anastrephae* showed clear and consistent EAD responses to three fruit volatiles emitted by SWD-infested strawberries. These were identified by GC-MS as ethyl butanoate, methyl butanoate and ethyl hexanoate (Figs. 1 and 2). These three compounds are common fruit esters frequently found during the ripening process of strawberry (Yan et al., 2018), and they are associated with ripening in several fruits. While they ubiquitous compounds, infestation by *D. suzukii* may accelerate the change in volatile profiles towards a riper blend, so the ability of parasitoids to sense these fruity esters suggests that the compounds act as general cues for suitable habitats to find their hosts.

**Figure 1.**
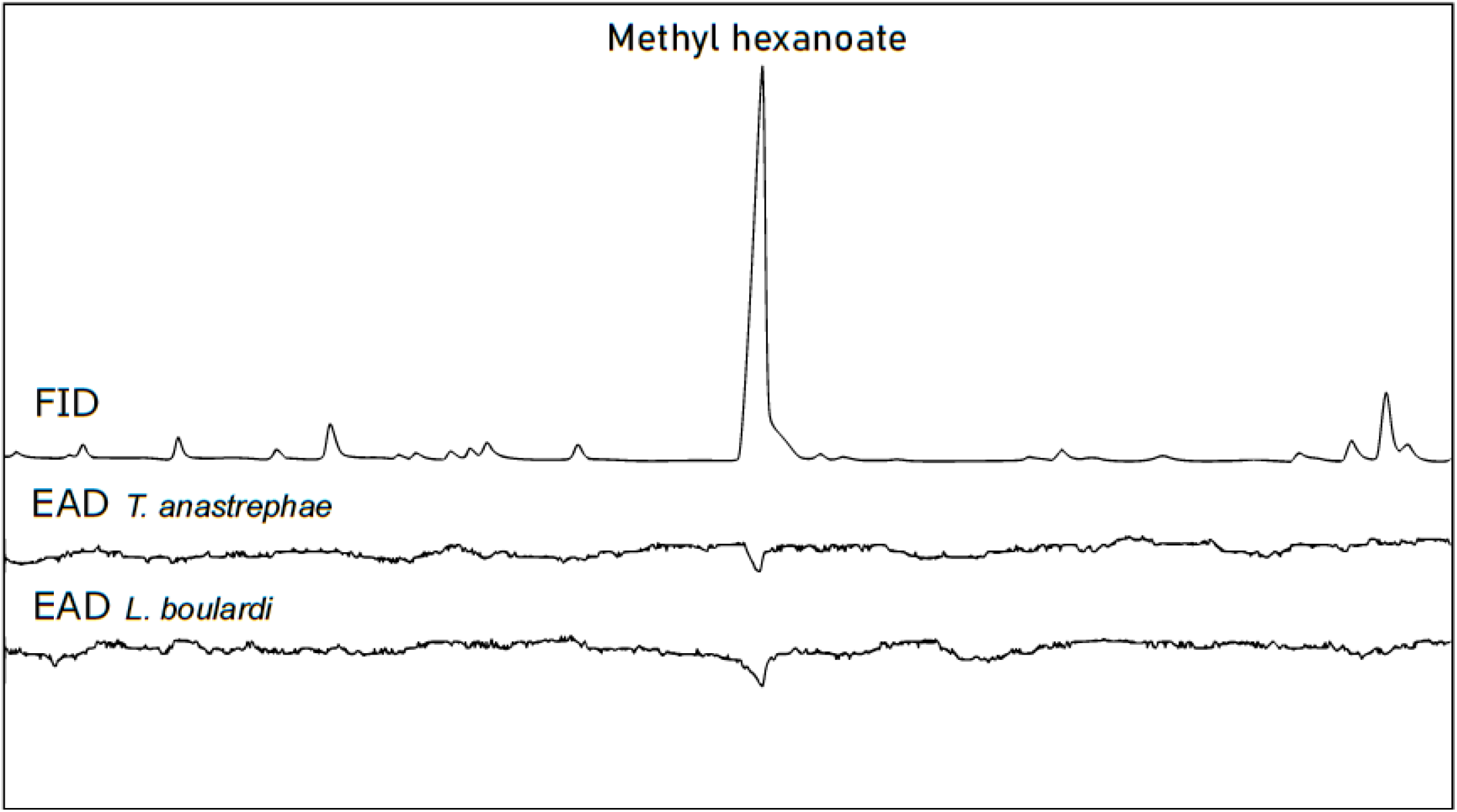
Antennal response of *T. anastrephae* and *L. boulardi* female wasps to VOCs of D. suzukii-infested strawberries five days post-infestation. First trace corresponds to FID signal and the following to *T. anastrephae* and *L. boulardi*, respectively. The main compound detected was identified by GC-MS as methyl hexanoate.

**Figure 2.**
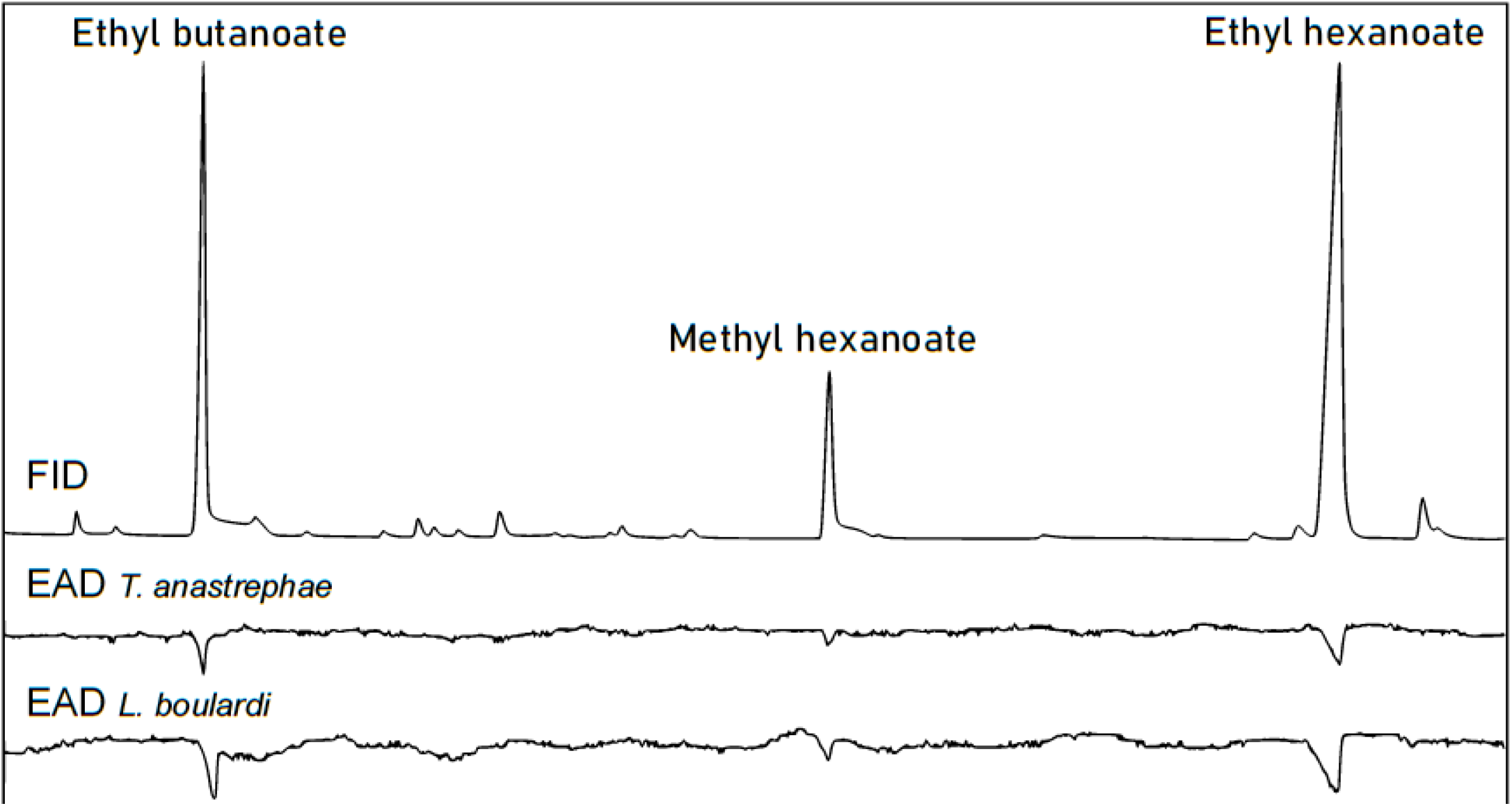
Antennal response of *T. anastrephae* and *L. boulardi* female wasps to VOCs of D. suzukii-infested strawberries ten days post-infestation. First trace corresponds to FID signal and the following to *T. anastrephae* and *L. boulardi*, respectively. The major compounds detected were identified by GC-MS as ethyl butanoate, methyl hexanoate and ethyl hexanoate (RT= 3.77, 6.28 y 8.32 min).

Interestingly, the relative composition of the fruit blends changed from the main methyl ester five days after SWD oviposition, to the ethyl esters later in the experiment. The antennae of both *L. boulardi* and *T. anastrephae* showed responses to all three esters, indicating that they are capable of detecting volatile cues associated with the early stages of SWD larval development. Indeed, five days after SWD oviposition methyl hexanoate triggered an EAD response (Fig. 1). Later on, ten days after oviposition, the relative amounts of both ethyl esters increased, triggering EAD responses even more pronounced that the response to methyl hexanoate (Fig. 2). As a whole, our results show that both parasitoids are capable of detecting infested fruit at different times during host development, with implications for biological control potential.

Our study with *L. boulardi* and *T. anastrephae* female antennae represent the first GC-EAD approach to understanding which chemical stimuli, within a complex fruit VOC blend, are drosophilid parasitoids cueing on for finding their prey. The detection of ethyl butanoate, methyl hexanoate and ethyl hexanoate esters agree with similar approaches conducted with *D. suzukii* itself and other *Drosophila* species (Stensmyr et al., 2003; Keesey et al., 2015; Revadi et al., 2015). In this sense, natural enemies exploit plant-related chemical cues similar to those used by herbivorous host species, due to the evolutionary closeness of the trophic interaction (Đurović et al., 2021, Yang et al., 2022).

Among the multiple approaches investigated for SWD management, biological control with parasitoids has been extensively studied and it has consolidated in recent years (Lee et al., 2019). In fact, inundative biological control is currently applied against *D. suzukii* in Europe, where *Trichopria drosophilae* (Hymenoptera: Diapriidae) is the most promising and commercially available biocontroler (Gonzalez-Cabrera et al., 2019; Rossi-Stacconi et al., 2018). In the United States, as a result of quarantine studies in Switzerland and California (US), and given the specificity shown by the *Ganaspis brasiliensis* (Hymenoptera: Figitidae) groups, a petition submitted to USDA-APHIS was approved for the release of *G. brasiliensis* G1 group (Beers et al., 2022). *Trichopria anastrephae* has been proposed in Brazil as useful in greenhouses (Vieira et al., 2020). Finally, larval parasitoids such as *Leptopilina* spp., which did not perform well in *D. suzukii*, can still reduce the hatching of adults and thus contribute to its mitigation (Knoll et al., 2017).

Beyond behavioural and performance studies, there is scant information regarding chemical stimuli used by drosophilid parasitoids. Most *Drosophila* species are not of agronomic concern, possibly explaining the somewhat delayed research on the sensory ecology of their parasitoids. The recent irruption of *D. suzukii* has placed them on the spotlight, and studies from different angles of parasitoid biology and ecology have become more common and will continue to grow, in view of developing efficient biological control tools. Foraging behaviour largely defines the beneficial impact of natural enemies, and their applicability should be evaluated in a tritrophic context. Techniques such as EAG and/or GC-EAD identify which semiochemicals modulate and establish tritrophic systems in nature. In a broader view, semiochemicals may enhance parasitoid attraction through odours to preserve their presence in the agroecosystem, or they can be used to train generalist wasps and thus optimize the encounter of hosts. Conversely, identifying non-detected semiochemicals may contribute to selecting volatiles for trapping systems, without affecting the parasitoid populations.

